# cycleX: multi-dimensional pseudotime reveals cell cycle and differentiation relationship of dendritic cell progenitors

**DOI:** 10.1101/222372

**Authors:** Yong Kee Tan, Xiaomeng Zhang, Jinmiao Chen

## Abstract

Advances in single-cell RNA-sequencing have helped reveal the previously underappreciated level of cellular heterogeneity present during cellular differentiation. A static snapshot of single-cell transcriptomes provides a good representation of the various stages of differentiation as differentiation is rarely synchronized between cells. Data from numerous single-cell analyses has suggested that cellular differentiation and development can be conceptualized as continuous processes. Consequently, computational algorithms have been developed to infer pseudotimes and re-ordered cells along developmental trajectories. However, existing pseudotime inference methods generate one-dimensional pseudotime in an unsupervised manner, which is inadequate to elucidate the effects of individual biological processes such as cell cycle and differentiation and the links between them. Here we present a method called cycleX which infers multi-dimensional pseudotimes to reveal putative relationship between cell cycle and differentiation during dendritic cell development. cycleX can be also applied to generate multi-dimensional pseudotime for the relationship among cell cycle, differentiation, trafficking, activation, metabolism and etc.

## Introduction

Interest in whole transcriptome analysis of single cells using single-cell mRNA sequencing has been growing tremendously, as it is able to identify the diversity between individual cells with different expression profiles in a heterogenous population. (Shapiro E, 2013). However, understanding and visualising these high dimensional datasets poses a major analytical challenge. Methods such as Principal Component Analysis (PCA) (Jolliffe, 1986), t-distributed stochastic neighbour embedding(t-SNE) (Maaten and Hinton, 2008) and Gaussian Process Latent Variable Model (GPLVM) (Lawrence, 2004) have been used for dimension reduction and visualisation of single-cell mRNAseq data. (Rostom et al., 2017). In addition to heterogeneity, methods such as Monocle2, Mpath, DPT and GPfates have been developed for pseudotime inference and trajectory construction. However, by using most existing methods the reduced dimensionalities or inferred pseudotimes represent a mixed effect of various biological processes such as cellular differentiation, cell cycle, activation, metabolism and etc. The relationship between the various biological processes is thus invisible.

Dendritic cells (DCs) are antigen-presenting cells which are vital for mounting an adaptive immune response to foreign antigens. A class of dendritic cells called conventional dendritic cells (cDCs) which refers to all DCs other than plasmacytoid DCs (pDCs). These cells populate most lymphoid and nonlymphoid tissues and function to sense tissue injuries, environmental and cell-associated antigens, and process and present phagocytosed antigens to T lymphocytes. (Merad et al., 2013). It is crucial to fully understand the development of cDCs for cDC-targeted immunotherapeutics. cDCs are known to differentiate from macrophage DC progenitor (MDP) in the bone marrow. The MDP give rise to both monocytes and to common DC progenitors (CDP). These CDPs are in turn able to give rise to precursors of classical dendritic cells (Pre-DC) which will further differentiate into conventional dendritic cells as cDC1 or cDC2 (Geissmann et al., 2010). Recently, single cell mRNA sequencing has been used to describe the development of individual precursor dendritic cells from its least differentiated state to the most differentiated state. (Schlitzer et al., 2015)

The same study has also revealed a connection between cell proliferation and differentiation in these DC progenitors. (Schlitzer et al., 2015) Genes encoding products associated to cell cycle always preceded gene expression that drove dendritic cell differentiation. (Schlitzer et al., 2015) Therefore in this manuscript, we explore the relationship of categories of genes, particularly differentiation and cell-cycle in one set of single-cell mRNA sequencing data of precursor dendritic cells using t-SNE and GPLVM by an analysis workflow that is implemented called ‘CycleX’. The inspiration of this methodology was born from the One-SENSE (Cheng et al., 2015) method where predefined categories of protein markers expression compared by dimension reduction to one dimension using t-SNE.

## Results

### t-SNE failed to visualise the relationship between differentiation and cell cycle

t-SNE is a dimension reduction technique that is commonly used for embedding highdimensional data into lower dimensional space of two or three dimension. The results generated in Fig. 1 to Fig. 3 uses the Barnes-Hut implementation of the algorithm available on R in package ‘Rtsne’ by using default values, of perplexity 30 for all sets of dimension reduction.

**Fig. 1:**
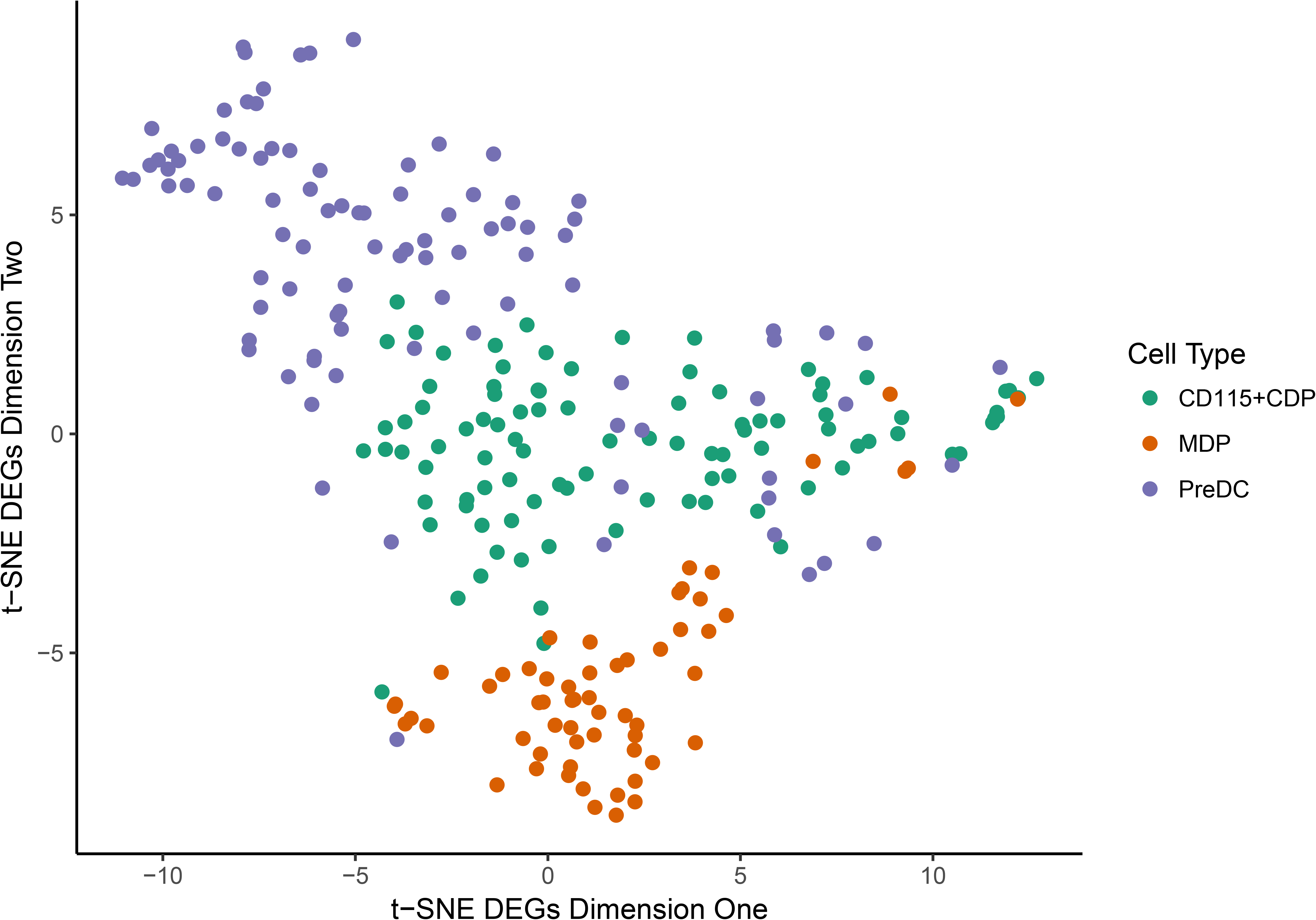
A 2D plot depicting t-SNE of 1783 differentially expressed genes reduced to two dimensions. X axis: t-SNE component 1, Y axis: t-SNE component 2. The cells are generally clustered by their cell type, which shows that t-SNE is able to group cells that are similar together using the differentially expressed genes obtained from the Seurat analysis. It is also able to separate them by their stage in differentiation. MDPs are labelled green, CD115+CDPs are labelled red and PreDCs are labelled blue.

**Fig. 2:**
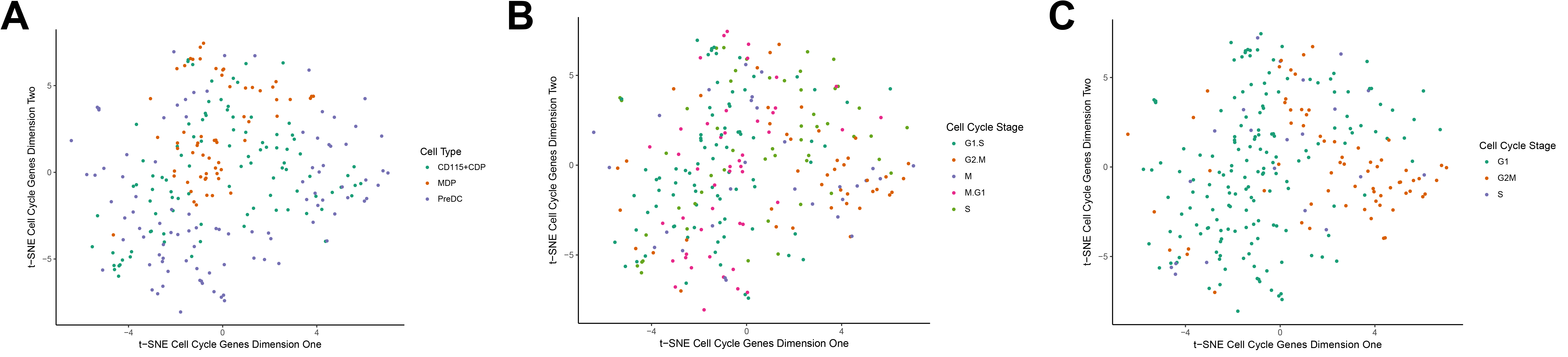
(**A**) A 2D plot of 1571 cell cycle genes obtained from AmiGO 2, reduced to two dimensions by t-SNE. Each point is a cell in 2D space. X axis: t-SNE component one Y axis: t-SNE component two. t-SNE is unable to separate the cells by their cell type, which is expected because using cell cycle genes do not differentiate them by their cell type. (**B**): A 2D plot of 1571 cell cycle genes obtained from AmiGO 2, reduced to two dimensions. Each point is a cell in 2D space. X axis: t-SNE component one Y axis: t-SNE component two. Cells are labelled by their Cell cycle stage according to Drop-Seq assignment algorithm (see Materials and Methods). (**C**): A 2D plot of 1571 cell cycle genes obtained from AmiGO 2, reduced to two dimensions by t-SNE. Each point is a cell in 2D space. X axis: t-SNE component one Y axis: t-SNE component two. Cells are labelled by their Cell cycle stage according to Cyclone, a computational method of assigning cell cycle stage (see Materials and Methods).

**Fig. 3:**
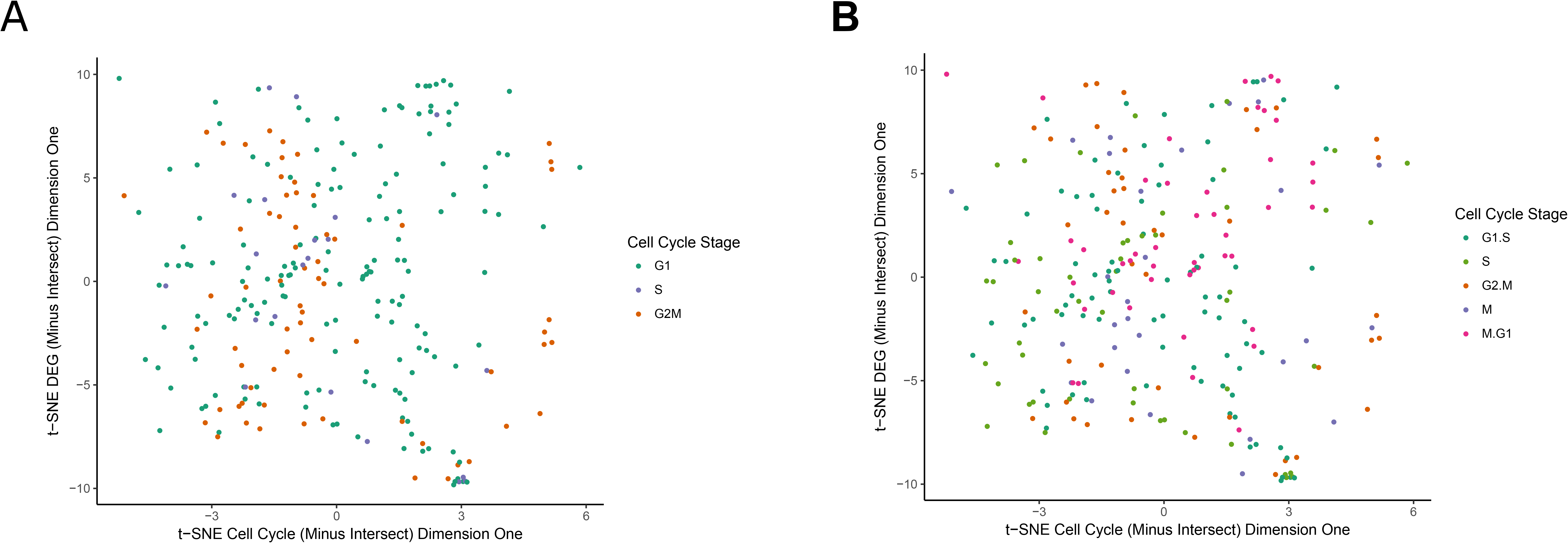
(**A**) 2D plot of t-SNE on both DEGs and Cell Cycle genes, reduced to one dimension. X axis: Cell Cycle genes reduced to one dimension by t-SNE. Y axis: DEGs reduced to one dimension by t-SNE. The cells are labelled to their cell cycle stage (G1, S, G2/M) according to ‘cyclone’. (**B**) 2D plot of t-SNE on both DEGs and Cell Cycle genes, each reduced to one dimension. X axis: Cell Cycle genes reduced to one dimension. Y axis: DEGs reduced to one dimension. The cells are labelled to their cell cycle stage (G1.S, S, G2.M, M, M.G1) according to Drop-Seq cell cycle assignment algorithm.

While t-SNE is able to resolve the cells by cell types in Fig. 1 using differentially expressed genes, it performed poorly when trying to resolve cells using cell cycle genes in Fig. 1A as it is unable to separate the CDPs and PreDCs, with the exception of clustering MDPs together. Furthermore, it is hard to perceive any form of order of cell cycle by using t-SNE to reduce cell cycle genes extracted from the original dataset. As seen from Fig. 2B and 2C, it is hard to assign any form of direction for cell cycle progression. In Fig. 2C although t-SNE is able to separate cell cycles staged in G2M closely together, G1 and S staged cells are mapped without any visible order. It is even harder to decipher any form of cell cycle progression from Fig. 2B.

By applying the methodology of One-SENSE (Cheng et al., 2016) which involves taking categories of gene sets and reducing each set to one dimension, one would expect to see a relationship between differentiation and cell cycle. Another thing to note is the removal of gene intersection between the two groups of gene sets, where there were 199 genes common between the differentially expressed genes and cell cycle genes. However, the cell cycle genes reduced to one dimension does not order the cells by their cell cycle stage assignment in Fig. 3A and Fig. 3B, even though cell cycle genes were used for the dimension reduction. While t-SNE is able to map cells in in Fig. 3A of stage G2M close together, this does not reveal any biological meaningful information between differentiation and cell cycle of these cells. There are also several disadvantages of using t-SNE (Wattenberg, et al., 2016), which makes t-SNE visualisations difficult to interpret. Firstly, the t-SNE algorithm doesn’t always produce the same results using the same inputs. Secondly, the perplexity value which balances the tendency to preserve local structure and global structure will affect the cluster shape and sizes of t-SNE outputs. Perhaps by running multiple plots at various perplexity, the spatial relationship of cells differentiating to its cell cycle could be clearer.

### Rational for using cycleX

cycleX is a novel algorithm that infers multi-dimensional pseudotimes by integrating GPLVM analysis using multiple categories of genes. GPLVM allows gene expression to follow a smooth (nonlinear) function over time (Rostom et al., 2017). There has been previous studies that used GPLVM to model the order of haematopoietic cells differentiation in a continuous spectrum (Macaulay, et al., 2016). The GPfates model also uses GPLVM to reconstruct CD4+ T cells differentiation fates into Th1/Tfh. (Lőnnberg et al., 2017) GPLVM using DEGs is able to cluster cells by cell type quite clearly as the clusters are visually distinct and separated (Figure 4). Component one of GPLVM is able to describe the differentiation path of the precursor dendritic cells from least differentiated to most differentiated in the correct order (Fig. 4). GPLVM using cell cycle genes only is also able to cluster MDPs, CDPs and PreDCs into their individual bands. However, there is a strong overlap between CDPs and PreDCs showing that GPLVM is still unable to resolve the class differences clearly, which is expected as the dimension reduction was done using only cell cycle genes. However, note that within the 1571 cell cycle genes, 199 of them have been identified by Seurat as differentially expressed genes to distinguish the cell types. These 199 genes are not excluded in this analysis while using GPLVM. This could have influenced GPLVM to cluster by cell type with better performance than t-SNE. A fair comparison would be to remove the DEGs identified by Seurat and see how well the cell types are clustered. Using GPLVM for dimension reduction allowed the visualisation of the progression of cell cycle in the diagonal as described in Fig. 6A and 6B, and it also did better in showing the progression of cell cycle than t-SNE as shown in Fig. 2B and Fig 2C.

**Fig. 4:**
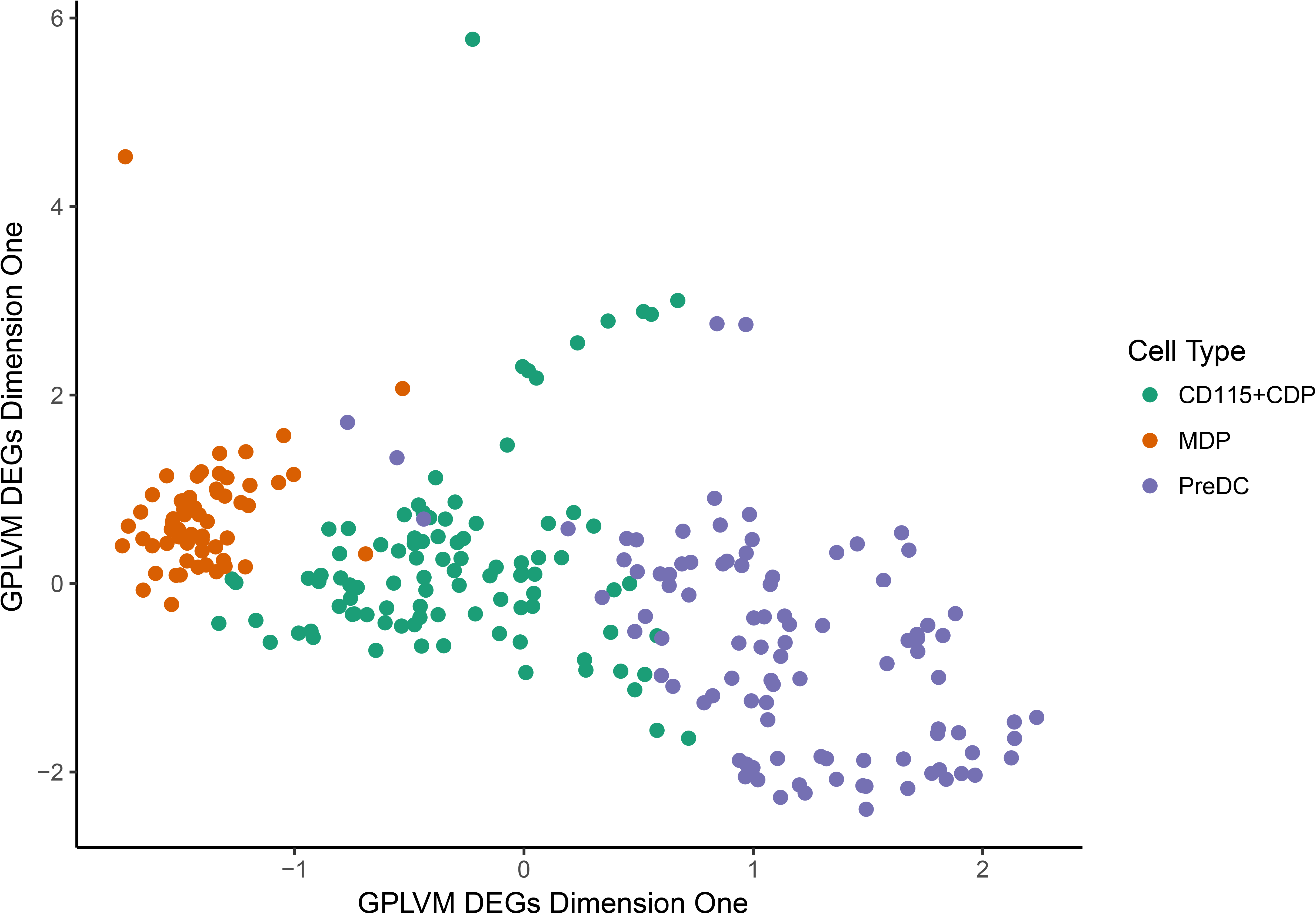
GPLVM dimension reduction of 1783 DEGs to two dimensions. X axis: GPLVM component 1. Y axis: GPLVM component 2. Cells are grouped by cell type.

**Fig. 5:**
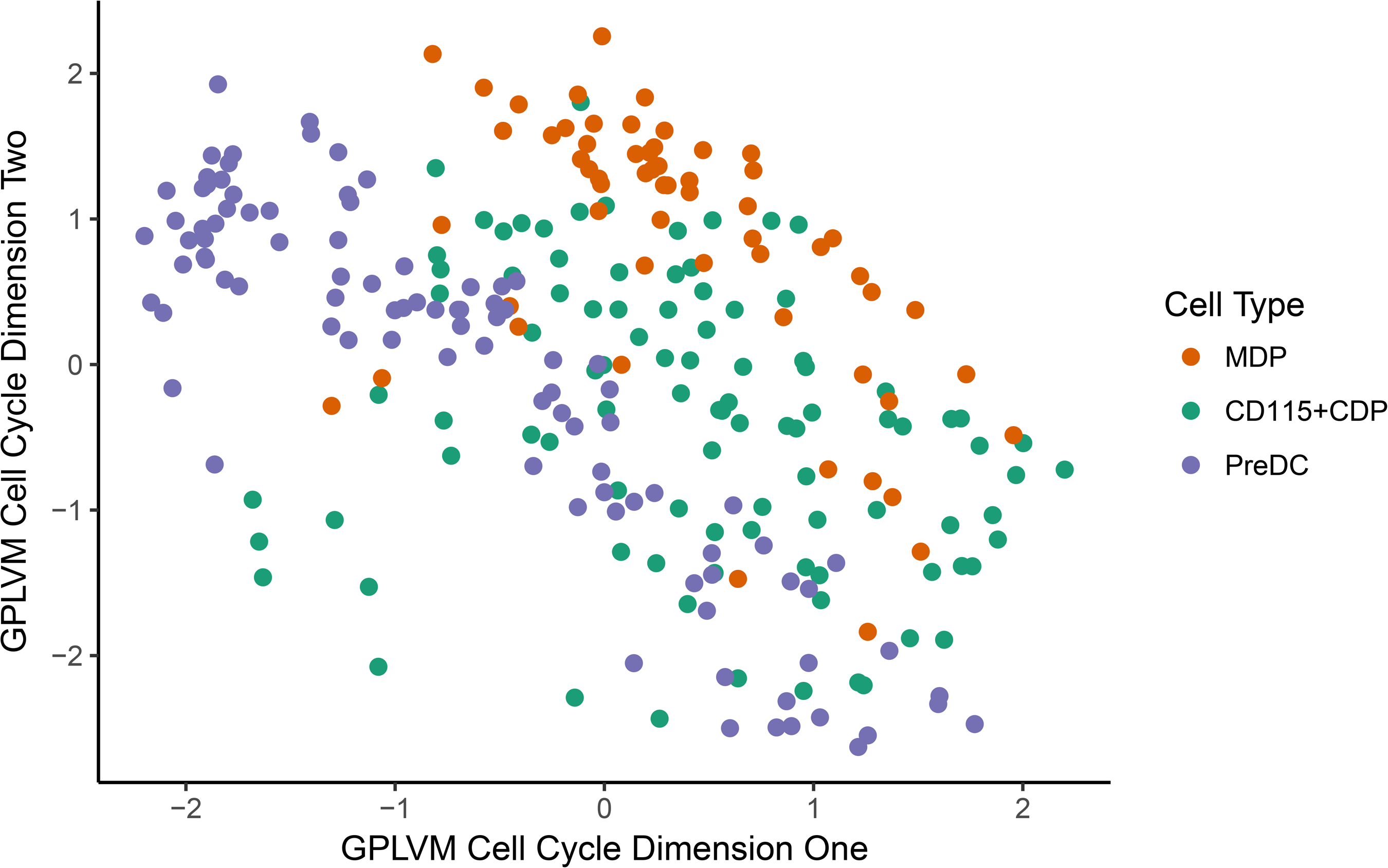
GPLVM dimension reduction of 1571 Cell cycle genes from AmiGO 2 gene ontology term to two dimensions. X axis: GPLVM component 1. Y axis: GPLVM component 2.

**Fig. 6:**
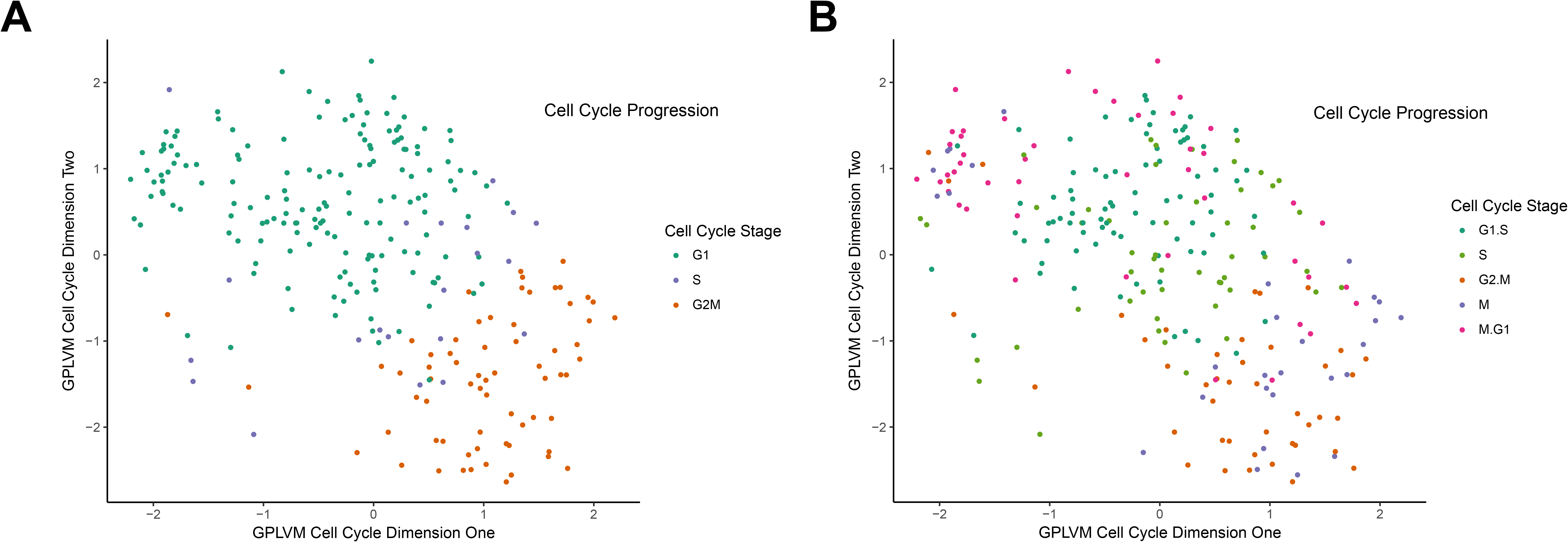
(**A**) Cell cycle progression (indicated by arrow) of 251 dendritic cells captured by GPLVM in component one and component two, cells are computationally assigned by ‘cyclone’. (**B**) Cell cycle progression (indicated by arrow) of 251 dendritic cells captured by GPLVM in component one and component two, cells are computationally assigned by Drop-Seq cell cycle algorithm. This figure also reveals the cell cycle’s circular nature. From G1 to S to G2 to M and back to G1.

The most interesting part about plotting GPLVM using two categories of gene sets, when differentially expressed genes are reduced to one dimension and cell cycle genes are reduced to one dimension and plotted against one another, an ‘X’ shape appears. (see Fig 7.) Even though we are not sure why this shape is the result of plotting two sets of high dimensional data together, biologically meaningful relationships between cell cycle and differentiation remains to be discovered. Between MDP and CDP, perhaps cells transition from one differentiation state to another when they are at the ‘S’ phase (see Fig. 7A and 7B). As cell cycle is a cyclic process, we fold the pseudotime along cell cycle dimension and generate 3D plot as shown in Figure 7C and 7D.

**Fig. 7:**
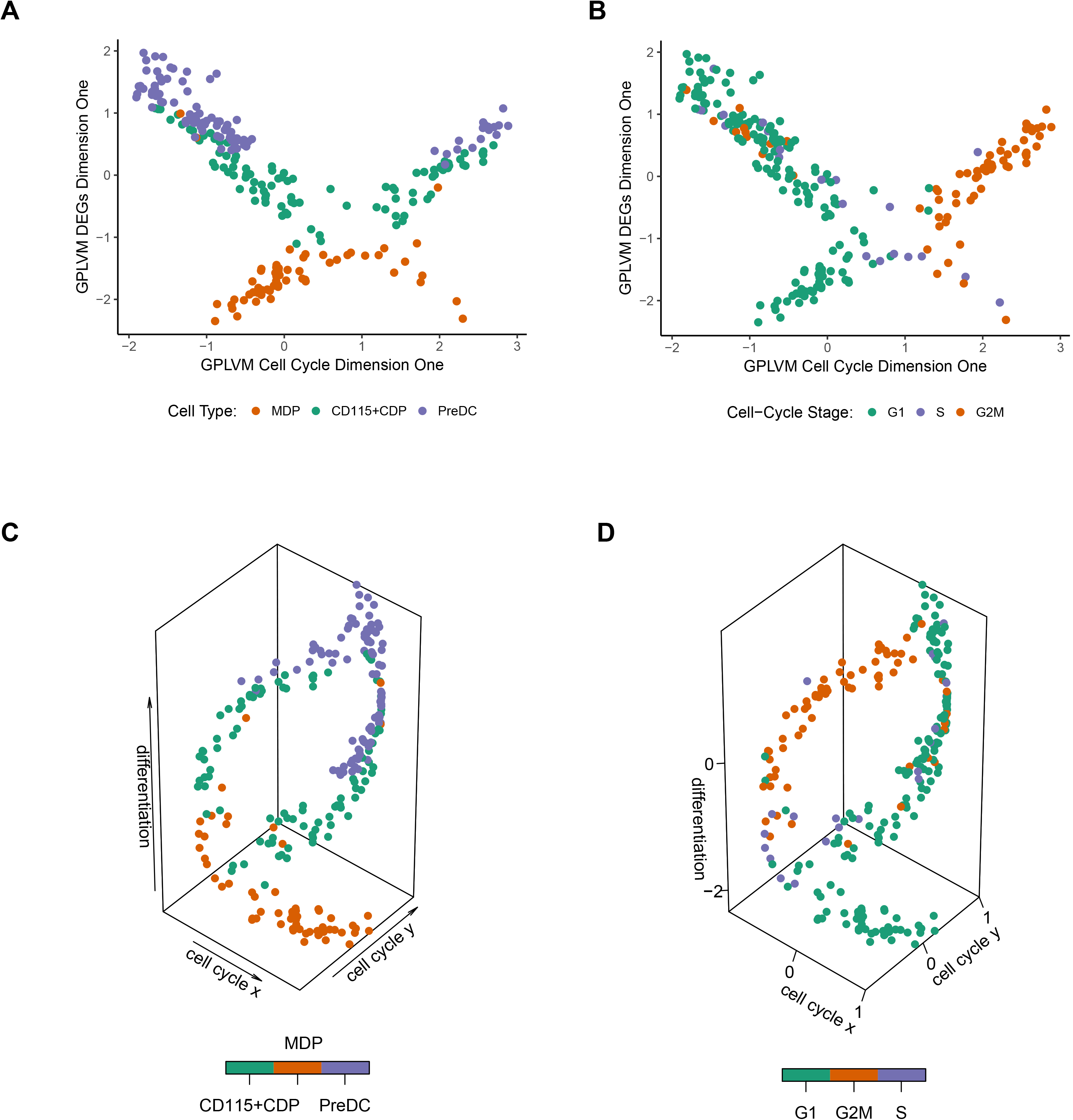
(**A**) 2D plot of 1374 differentially expressed genes vs cell cycle genes. The X axis represents the cell cycle genes reduced to one dimension by GPLVM. The Y axis represents the DEGs reduced to one dimension by GPLVM. The cell types (MDP, CDP, PreDC) are labelled in colour. (**B**) 2D plot of 1374 differentially expressed genes vs cell cycle genes. The X axis represents the cell cycle genes reduced to one dimension by GPLVM. The Y axis represents the DEGs reduced to one dimension by GPLVM. The cell cycle stages (G1, S, G2M) are assigned to each cell, labelled in colour by a cell cycle assignment function called ‘cyclone’. (**C**) 3D plot of 1374 differentially expressed genes vs cell cycle genes. The X axis represents the cell cycle genes reduced to one dimension by GPLVM. The Y axis represents the DEGs reduced to one dimension by GPLVM. The cell types (MDP, CDP, PreDC) are labelled in colour. (**D**) 3D plot of 1374 differentially expressed genes vs cell cycle genes. The X axis represents the cell cycle genes reduced to one dimension by GPLVM. The Y axis represents the DEGs reduced to one dimension by GPLVM. The cell cycle stages (G1, S, G2M) are assigned to each cell, labelled in colour by a cell cycle assignment function called ‘cyclone’.

In order to visualise the mapping of cell cycle with differentiation, we assigned the 251 individual dendritic cells to a cell cycle stage. Using ‘cyclone’ method, from the Scialdone et al. (2015) paper, all cells were assigned stages to either ‘G1’, ‘S’ or ‘G2M’. However, with no methods to measure the accuracy of the assignment, only comparisons with another method can be made. Another method to assign cell cycle stage was available from Macosko et al., (2015) where cells were assigned to any of the 5 stages ‘G1.S’, ‘S’, ‘G2.M’, ‘M’ or ‘M.G1’. From a visualisation standpoint of view (Fig 7B and supplementary Fig1), having three classes of cell cycle stage makes it easier to notice changes as compared to having five classes of cell cycle stage assignment. This is especially more apparent in Fig 7B where the G1 stage cells are on the left of the plot, and G2M staged cells on the right with cells in the S phase being in the centre of the ‘X’. Comparing Fig 7B and supplementary Fig1 also shows that both methods agree with one another where cells labelled G1 in Fig. 7B corresponds to cells that are labelled G1.S and M.G1 as both are on the left side of the 2D plot. Also, cells labelled G2M in supplementary Fig. 1 corresponds to cells labelled G2.M and M in Fig. 7B on the right side of the 2D plot.

cycleX is also robust to the removal of cells, when certain group of cells are removed (MDP/CDP/PreDC) from the original dataset, the shape of X still remains (see Fig. 8A-8C). Furthermore, removing 10 or 50 random cells also results in a similar shaped plot (see Fig. 8D-8E).

**Fig. 8:**
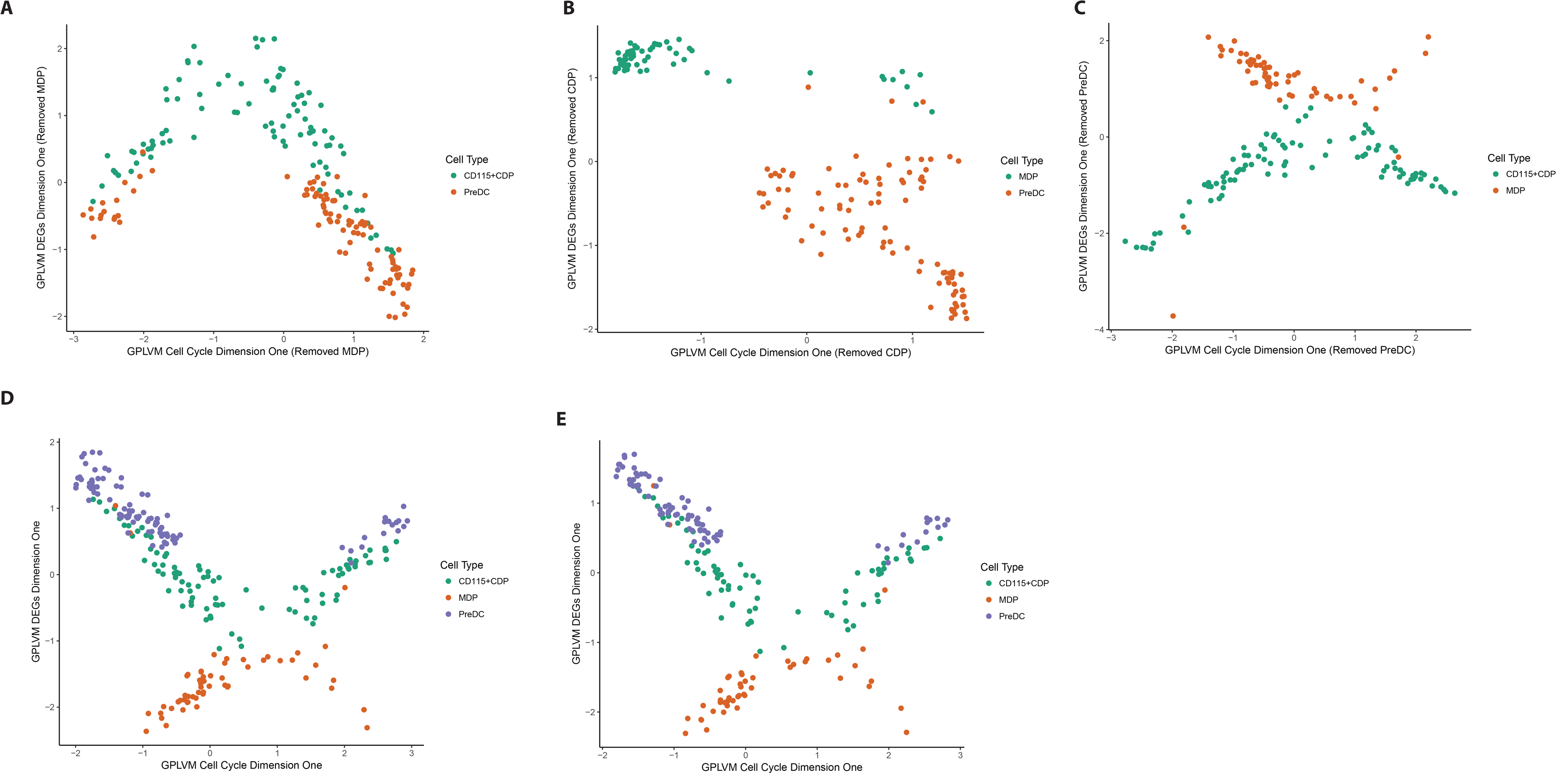
2D plots of differentially expressed genes vs cell cycle genes. MDPs are removed from the dimension reduction using GPLVM. (**A**) MDPs removed (**B**) CDPs removed (**C**) PreDCs are removed. (**D**) 10 Random cells removed (E) 50 Random Cells removed

## Discussion

Single Cell mRNA sequencing data is able to reveal intimate relationships such as the differentiation of cells from least differentiated to most differentiated to even displaying how their cell cycle progresses. These are revealed with dimensional reduction methods such as t-SNE and GPLVM, with the help of gene set annotation to visualise cell to cell heterogeneity. GPLVM has been shown to be able separate cells based on their cell type and even having the potential to reveal cell cycle effects. cycleX based on GPLVM dimension reduction has been shown to have the capability to order cells based on their cell cycle progression and visualize the relationship between cell differentiation and cell cycle. Since cell cycle effects are cycling, GPLVM is able to model the cycling effect with a smooth function similar to a sine curve due to the usage of a Gaussian kernel. An alternative technique, Bayesian GPLVM has been used to depict ‘pseudo time’ of cell differentiation. (Macaulay et al., 2016) One way to improve the visualisation of the method is to order the cells by their ‘pseudo time’ of differentiation and cell cycle progression, and look at the points in time where the cell cycle and the transition of MDP to CDP, CDP to PreDCs occur. Using this method we can find out at which cell cycle stage the cells will differentiate.

Knowing the cell cycle stage of the cell is important as cell cycle effects have been known to confound global gene expression (Buettner et al., 2015). There are only two known methods of assignment cell cycle stage so far which are ‘cyclone’ and the method presented in Macosko et al., (2015). Furthermore, the cell cycle does not only consist of cells from G1.S to M.G1 phase, it also includes the G0 phase, where we know that cells do not actively divide at this stage. (Scialdone et al., 2015; Macosko et al., 2015) Standardised methods of cell cycle assignment for single cell mRNA sequencing data should be introduced in the future to allow accurate assignment of cell cycle stage. This will allow us to gain further insight into the relationship of cell cycle vs differentiation.

The question remains if cell cycle and differentiation have an intimate relationship as the previous study has revealed. There has been evidence that cell fate decisions are cell cycle dependent (Pauklin and Vallier, 2013). Perhaps future work should explore on utilising CycleX to compare different gene sets available and other single cell mRNA sequencing datasets, to further investigate the relationship between cell cycle and differentiation. Moreover, cycleX can also be used to reveal other intimate relationships such as between cell cycle, metabolism, activation, trafficking or other gene sets of interest.

## Materials & Methods

Most of the analysis was done using R environment, with RStudio. Only one program required Python and was implemented on Jupyter notebook.

### Dataset

A .txt file (26.6MB) containing transcript per million reads (TPM) of 43280 genes measured in 251 precursor dendritic cells from a previous study (Schlizter et al., 2015) were used in this analysis, generated from scRNAseq. RSEM was used to get the alignment and TPM values. (Li and Dewey, 2011) Another .txt file (4KB) obtained from the previous study containing each cell’s sample identification and group identification was used to label the cells during differentiation.

### Analysis and workflow of CycleX

The workfow of CycleX is illustrated in Fig 9. Cell cycle genes and differentially expressed genes (DEGs) were extracted and GPLVM was applied to each gene set to reduce dimensionality to one. Subsequently GPLVM dimension one using DEGs was plot against GPLVM dimension one using cell cycle genes. As cell cycle is a cyclic process, we fold the x-axis (i.e. the cell cycle pseudotime) and generated 3D cycleX plot using

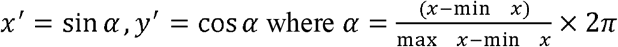

**Fig. 9:**
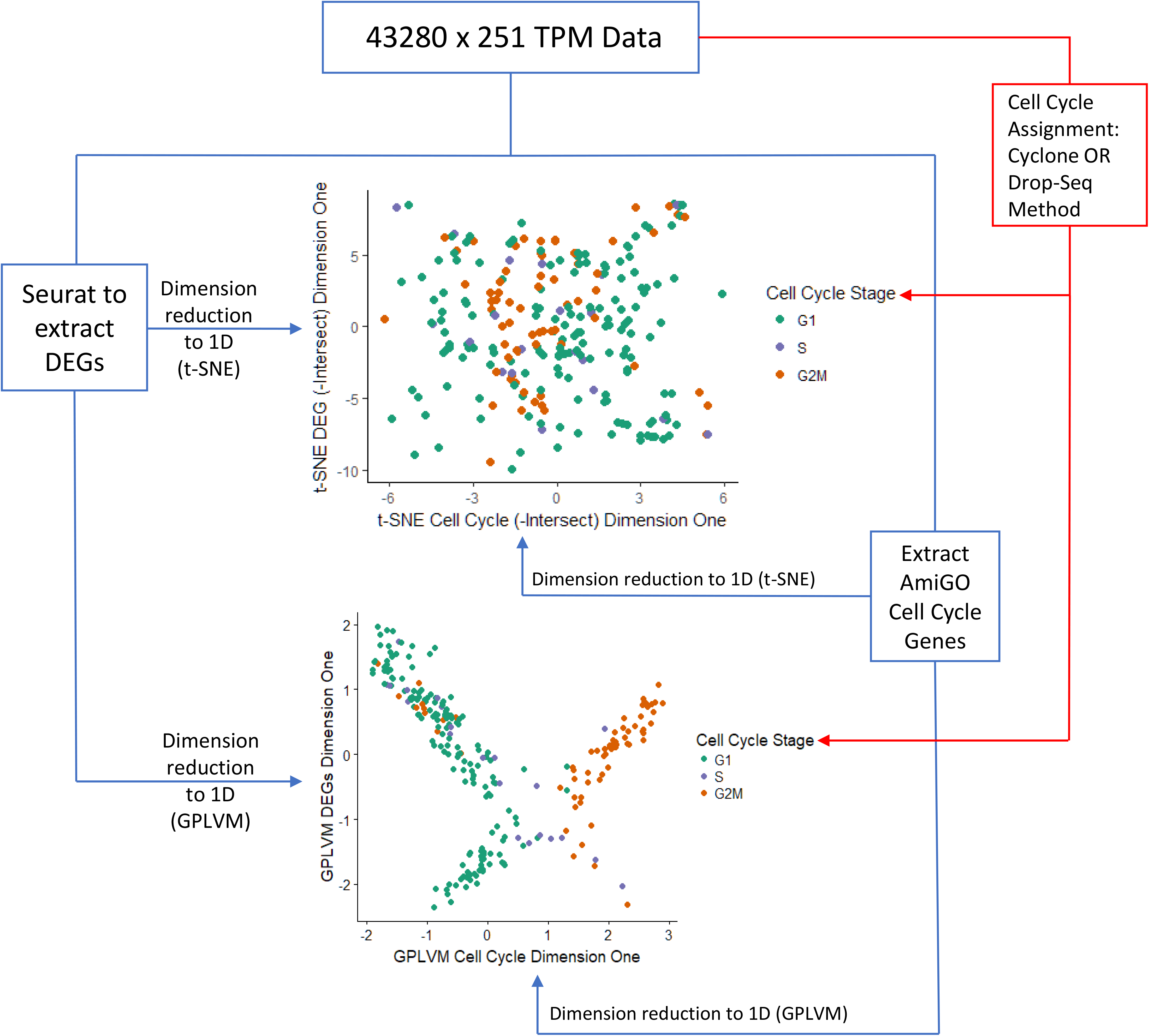
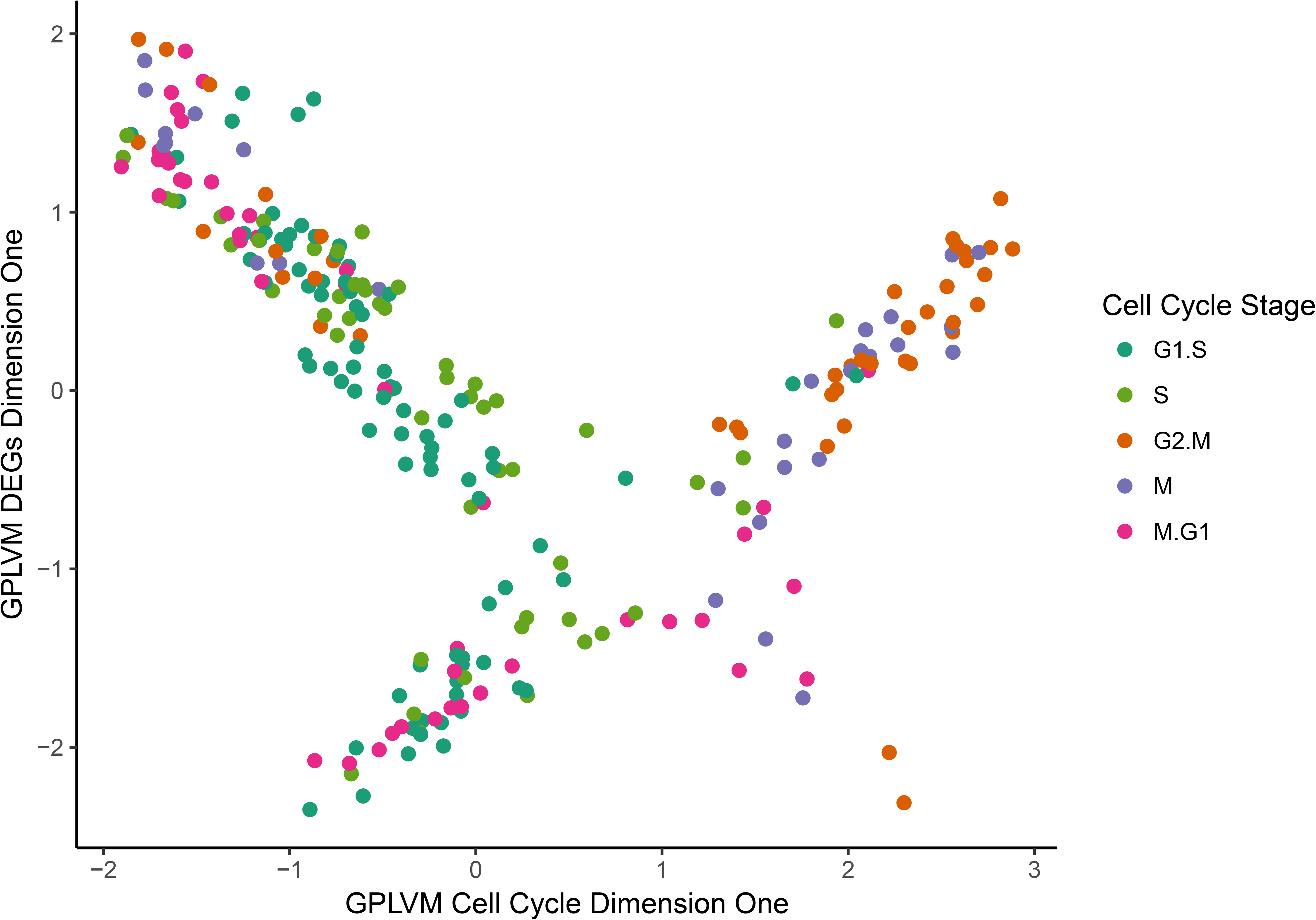
Workflow of CycleX to explore relationship between cell cycle and differentiation in single cell mRNA sequencing data.

### Seurat to obtain differentially expressed genes for dendritic cell differentiation

The file is loaded to an R environment for analysis. To obtain the differentially expressed genes (DEGs) among the three cell types (MDP, CDP, PreDC) an R package called ‘Seurat’ version 2.1.0 (http://satijalab.org/seurat/install.html) along with two dependant packages ‘dplyr’ and ‘Matrix’ were used. All of these packages were downloaded from CRAN (https://cran.r-project.org/) using RStudio. One interesting thing to note is that it is important to keep track of package versions when using them as the functions used may be modified without the user knowing, resulting in inconsistencies in result output. Between version 2.0.1 and 2.1.0 of Seurat, the function ‘FindAllMarkers’ resulted in different number of DEGs. Version 2.0.1 resulted in 1375 DEGs while version 2.1.0 resulted in 1783 DEGs of which 1375 DEGs are a subset of the 1783 DEGs found in the later version.

The dataset was first log normalized with a scale factor of 10,000. A function ‘ScaleData’ scales and centres genes in the dataset causing effects such as ‘nUMI’ and mitochondrial gene expression to be regressed out. Differentially expressed genes (DEGs) based on the cell groups are then found using the function ‘FindAllMarkers’, and 1784 DEGs were found. This modified dataset will be used as described later.

### Obtaining Cell Cycle Genes

A list of 1621 cell cycle annotated genes were obtained from AmiGO 2. (http://amigo.geneontology.org/amigo/search/bioentity?q=*:*&fq=regulates_closure:%22GO:0007049%22&sfq=document_category:%22bioentity%22) The search was further refined by selecting for *Mus Musculus* under type of organism.

### Extracting Cell Cycle Genes from dataset

Cell cycle genes and their expression values were extracted from the original dataset. Out of 1621 genes from the GO term, 1571 genes were found to be in the dataset. I have written a short script for extracting cell cycle genes from the original data set using R. This modified dataset will be used as described later.

### Computational Assignment of Cell Cycle Stage

Two methods were used to assign the cell cycle phases for each cell. The first method was a classification algorithm based on the idea of relative expression of ‘marker pairs’. This is done by using the R function ‘cyclone’ in the ‘scran’ package (http://bioconductor.org/packages/scran/) available on Bioconductor with the aid of R package ‘biomaRt’ (http://bioconductor.org/packages/biomaRt/) which is also available on Bioconductor. To assign the cell cycle stage for each of the 251 precursor dendritic cells, the Mouse Genome Informatics (MGI) gene symbols in the original dataset are converted into Ensembl Gene Ids using the function ‘getBM’ from the ‘biomaRt’ package. Using the dataset with converted gene ids as an input in ‘cyclone’, the cell cycle stage is assigned to each cell, using the training data available from the package.

The second method of assigning cell cycle stage dubbed Drop-Seq in this report references the cell-cycle analysis of HEK and 3T3 cells in (Macosko et al, 2015). Gene sets reflecting five phases of HeLa cell cycle (G1/S, S, G2/M, M, M/G1) taken from (Whitefield et al., 2002) refined by Macosko et al. (2015) were used to assign the cell cycle stage. Each cell has five scores for each of the five cell cycle stages. The assignment of the stage is based on the most positive score. A script was implemented to assign cell cycle stage in R environment.

### Dimension reduction for visualisation

#### t-SNE

The t-SNE algorithm (Maaten & Hinton, 2008) is available through a Barnes-Hut implementation on CRAN in the function ‘Rtsne’. The mathematical description of t-SNE can be found (https://lvdmaaten.github.io/tsne/). t-SNE was applied to two modified datasets, one with cell cycle genes only and the other modified dataset with DEGs. By following One-SENSE methodology, the intersect of genes between cell-cycle and DEGs were removed. The first set of gene belonging to DEGs is first reduced to two dimensions and the first two components plotted in 2D to visualise how well t-SNE is able to cluster the cells by cell type and observe their order of differentiation. The second set of genes belonging to cell cycle is then also reduced to two dimensions and the first two components plotting in 2D to visualise how well t-SNE is able to cluster the cells by cell type and also observe the progression of differentiation as well as progression of cell cycle. (See Fig.5 and 6) Next, using the same set two sets of genes, which are the cell cycle genes and DEGs, each of them were then reduced to only one dimension. DEGs reduced to one dimension was plotted as the Y axis and cell cycle genes reduced to one dimension was plotted on the X axis, resulting in a 2D plot of cells (see Fig. 3) The plot was made using ‘ggplot’ function from ‘ggplot2’ package available on R.

### Gaussian Process Latent Variable Model

Gaussian Process Latent Variable Model (GPLVM) (Lawrence, 2004) is implemented on Python in the ‘GPy’ package. (http://gpy.readthedocs.io/en/deploy/index.html) GPLVM was applied to the same two modified datasets, however, without removing the gene intersect between cell cycle genes and the DEGs. Both sets of genes were then reduced to 2 dimensions to see how well GPLVM is able to separate the cells based on their differentiation. Then, both the cell cycle genes and the DEGs are reduced to one dimension and their positions plotted against one another in a 2D plot using ‘ggplot’ function from ‘ggplot2’ package available on CRAN. The script for GPLVM dimension reduction is available in the format of a .ipynb file which was referenced from an online lab session. (http://gpss.cc/gpss13/assets/lab3.pdf)

